# The Moorfields AMD Database Report 2 - Fellow Eye Involvement with Neovascular Age-related Macular Degeneration

**DOI:** 10.1101/615252

**Authors:** Katrin Fasler, Gabriella Moraes, Dun Jack Fu, Siegfried K. Wagner, Eesha Gokhale, Karsten U. Kortuem, Reena Chopra, Livia Faes, Gabriella Preston, Nikolas Pontikos, Praveen J. Patel, Adnan Tufail, Aaron Y. Lee, Konstantinos Balaskas, Pearse A. Keane

**Affiliations:** NIHR Biomedical Research Centre at Moorfields Eye Hospital NHS Foundation Trust and UCL Institute of Ophthalmology, London, UK; Department of Ophthalmology, University Hospital Zurich, Zurich, Switzerland; University Eye Hospital, Munich, Germany; Eye Clinic, Cantonal Hospital of Lucerne, Lucerne, Switzerland; Department of Ophthalmology, University of Washington, Seattle, USA; School of Biological Sciences, University of Manchester, Manchester, UK

**Keywords:** Age-related macular degeneration, Choroidal neovascularization, Ranibizumab, Aflibercept, Anti-vascular endothelial growth factor, Fellow eye, Electronic medical record, Visual acuity

## Abstract

**Background/Aims:** Neovascular age-related macular degeneration (nAMD) is frequently bilateral, and previous reports on ‘fellow eyes’’ have assumed sequential treatment after a period of treatment of the first eye only. The aim of our study was to analyse baseline characteristics and visual acuity (VA) outcomes of fellow eye involvement with nAMD, specifically differentiating between sequential and non-sequential (due to macular scarring in the first eye) anti-vascular endothelial growth factor treatment and timelines for fellow eye involvement.

**Methods:** Retrospective, electronic medical record database study of the Moorfields AMD database of 8174 eyes/120,756 single entries with data extracted between October 21, 2008 and August 9, 2018. The dataset for analysis consisted of 1180 sequential, 413 nonsequential, and 1110 unilateral eyes.

**Results:** Mean VA of sequentially treated fellow eyes at baseline was significantly higher (62±13), VA gain over two years lower (0.65±14), and proportion of eyes with good VA (≥20/40 or 70 letters) higher (46%) than the respective first eyes (baseline VA 54±16, VA gain at two years 5.6±15, percentage of eyes with good VA 38%). Non-sequential fellow eyes showed baseline characteristics and VA outcomes similar to first eyes. Fellow eye involvement rate was 32% at two years, and median time interval to fellow eye involvement was 71 (IQR 27-147) weeks.

**Conclusion:** This reports shows sequentially treated nAMD fellow eyes have better baseline and final VA than non-sequentially treated eyes after 2 years of treatment. Sequentially treated eyes also had a greater proportion with good VA after 2 years of treatment.

**PRECIS:** Depending on age, fellow eye involvement occurs in 32% of patients with neovascular AMD by two years. Fellow eyes generally maintain better vision, except in cases where late-stage disease in the first eye was untreated.

## INTRODUCTION

Anti-vascular endothelial growth factor (VEGF) therapy has revolutionised the treatment of neovascular age-related macular degeneration (nAMD). Two anti-VEGF drugs, ranibizumab and aflibercept, form the mainstay of treatment and received approval from the Food and Drug Administration (FDA) in 2006 and 2011 respectively (bevacizumab is also widely used in an off-licence manner). The pivotal randomised controlled trials (RCTs) that led to the approval of these agents only included one eye per patient as a means of preventing bias due to correlation between eyes.[1,2] This is a necessary step in RCTs, as not accounting for this effect can lead to overestimation of precision and a falsely low p-value.[3] However, this systematically excludes eyes of patients who subsequently develop nAMD in their fellow eye. From a patient’s perspective however, vision-related quality of life does not only depend on the course of visual acuity (VA) in the first treated eye.[4] Legal requirements largely focus on the VA of the better seeing eye. For example, in the United Kingdom, the VA standard for driving is 20/40 and the limit for obtaining a certificate of severe sight impairment is 20/400 (tested binocularly or in the better seeing eye).[5,6] Additionally, patients with bilateral nAMD have functional impairments that lead to a high socioeconomic burden.[7–9]

Data on treatment of fellow eyes, specifically sequentially treated fellow eyes, have been reported in small retrospective studies and in one large multicentre electronic medical record (EMR) report.[10–13] These studies concluded that fellow eyes commenced treatment with a higher baseline VA in comparison to the first treated eyes. In addition, they had a smaller gain in VA over time due to the relatively higher baseline VA, i.e., a ceiling effect. However, these studies do not account for non-sequentially treated fellow eyes, i.e., eyes starting treatment for nAMD in fellow eyes with an untreated first eye (e.g. due to development of nAMD in the first eye before anti-VEGF approval or late presentation at first eye involvement). Given that involvement of the fellow eye has a substantial impact on vision-related quality of life, accounting for patients that already have poor vision from macular scarring in their first eye clearly is important.[14]

The electronic medical record (EMR) database at Moorfields Eye Hospital NHS Foundation Trust, London, United Kingdom (UK), represents an ideal source to explore the unanswered questions on fellow eye nAMD outcomes. [Fasler, One and Two Year Visual Outcomes from the Moorfields Age-related Macular Degeneration Database, submitted to BMJ open, 2019] This database consists of over 8000 treatment naïve eyes with over 120,000 single entries and has undergone extensive manual data cleaning. Key elements that distinguish its quality compared to others include the completeness of data due to the mandatory input of relevant fields such as VA, the consistency of VA measurements using Early Treatment Diabetic Retinopathy Study (ETDRS) letters, the lack of requirement to merge data from different sites and systems, the standardised treatment scheme following national guidelines, and the ability to directly access the raw imaging data from each patient visit.[15,16]

The aim of this study was to analyse baseline characteristics and VA outcomes of fellow eyes (sequentially and non-sequentially treated) undergoing anti-VEGF therapy for nAMD, as well as the timelines for fellow eye involvement. We compare fellow eye outcomes to those of the respective first eyes of sequentially treated fellow eyes.

## METHODS

### Study Population

Data for this retrospective, comparative, non-randomised cohort study was extracted from the Moorfields AMD Database, consisting of 8174 treatment-naïve eyes / 6664 patients with 120,756 single entries acquired between October 21, 2008 and August 9, 2018. This has been reported in detail elsewhere.[17]

The complete dataset for analysis of the current study consisted of the 3890 eyes / 2710 patients (Supplementary sFigure 1). Of these, 1180 patients had sequentially treated fellow eye involvement, while 413 eyes were non-sequentially treated fellow eyes (i.e., untreated macular scarring in their respective first eyes). The 1117 unilateral/singular eyes (only treated in one eye over the observed period without advanced AMD in their fellow eye) were used for the survival analysis of fellow-eye involvement. Definition of sequential involvement was a time interval of ≥28 days between the first injection of first and fellow eye over the course of the observed time period. The presence of a macular scar was manually graded in fundus photographs and optical coherence tomography (OCT) (Topcon 3D OCT, Topcon, Japan) scans. An exemplar case for each group is shown in Figure 1.

**Figure 1:**
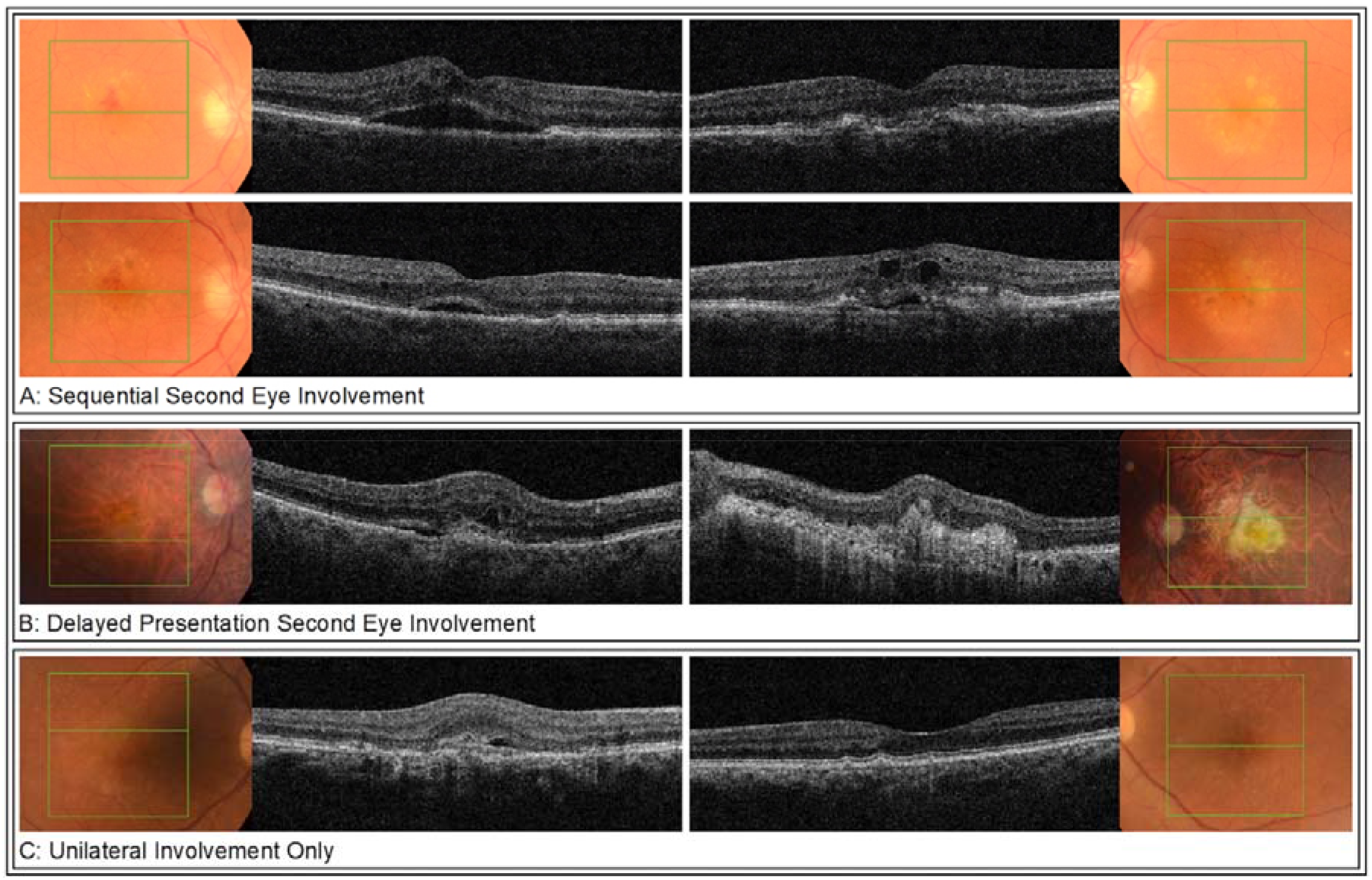
Representative fundus photographs and OCT-images for each of the groups at involvement of the respective eye(s). Sequential treatment fellow eye involvement (A) of a 83-year-old male. *Top row* shows findings at first eye involvement: neovascular AMD with intraretinal hemorrhage, intraretinal fluid, and subretinal fluid on OCT in the right eye, and early dry AMD in the left eye. One month later, new neovascular changes with intra- and subretinal fluid on OCT were found incidentally in the left eye, indicating second eye involvement (*bottom row*). Non-sequential treatment fellow eye involvement (B) of a 86-year-old male. Examination at first presentation revealed neovascular AMD in his right eye with intra- and subretinal fluid on OCT while in the left eye, a disciform scar with a visual acuity of hand movements was present. Unilateral involvement only (C) of a 86-year-old male. Examination at baseline showed neovascular AMD in the right eye with predominantly subretinal fluid on OCT and early AMD with drusen in the left eye. Until time of data extraction, there has been no progression to neovascular AMD in the left eye. OCT, optical coherence tomography; AMD, age related macular degeneration

Approval for data collection and analysis was obtained from the Institutional Review Board at Moorfields (ROAD17/031) and adhered to the tenets set forth in the Declaration of Helsinki.

### Efforts to Minimise Bias

To minimise survival bias/loss to follow-up (LTFU), all first and fellow eyes that did not complete follow-up were manually validated for the correct date of first injection. All unilateral eyes (1521) underwent manual verification for the presence of a macular scar secondary to end-stage AMD in the fellow eye. We chose not to substitute missing values, but clearly show results for cohorts that complete a certain follow-up period. Visual acuities below measurable ETDRS letters were converted to logMAR 2.0/-15 letters, logMAR 2.3/-30 letters and logMAR 2.7/-50 letters for count fingers, hand movements, and light perception respectively.[18]

### Outcome Measures

The primary outcomes were analogous to the pivotal RCTs, and as recommended by The International Consortium for Health Outcomes Measurement (ICHOM) AMD study group: mean change in VA from baseline as measured using ETDRS letters, proportion of eyes gaining ≥ 5 letters, proportion of eyes with stable vision (change in VA <15 letters to baseline), proportion of eyes with good vision (≥20/40 or 70 letters), and proportion of eyes with poor vision (≤20/200 or 35 letters).[1,2,19,20] Secondary outcomes included the number of injections and time to involvement of fellow eyes. Definitions for one-year and two-year outcome dates were taken from previous real-world studies as visits closest to 52 weeks and 104 weeks post baseline date within ±8 weeks.[21,22]

### Statistical Analysis

The data were analysed using the statistics software R (https://www.r-project.org/; provided in the public domain by R Core team 2017 R Foundation for Statistical Computing, Vienna, Austria). The ggplot2 package was used for plots. The eye was defined as unit of analysis. Descriptive statistics included mean +/-95% confidence interval (CI), and median, where appropriate. Differences between groups were evaluated using Mann Whitney U test and Pearson Chi-Square. A *p* value of < 0.05 was interpreted as statistically significant.

### Data Sharing Statement

De-personalised data as well as the code used for analysis for this study will be openly available from the Dryad Digital Repository https://doi.org/…. This should allow both for independent replication of our results as well as additional novel analyses. Depersonalisation was carried out through hash function anonymisation of patient identification numbers, and replacement of appointment dates with follow-up days to baseline. Approval of adequate depersonalisation was obtained by Moorfields Information Governance.

## RESULTS

### Baseline Characteristics

Of the 2710 patients starting treatment in one eye, 1180 (44%) developed fellow eye involvement, 413 (15%) were identified as non-sequentially treated fellow eye involvement, whereas 1117 (41%) were singular/unilateral eyes. Supplementary sFigure 1 shows the flow chart for eyes through the analysis. Mean baseline VA was 54±16 letters for first eyes and 62±13 letters for fellow eyes in sequentially treated patients, and 52±16 letters for non-sequentially treated fellow eyes (Table 1). In sequentially treated patients, fellow eyes had a significantly higher baseline VA than first eyes (p <0.001); more than 40% of fellow eyes had a VA of ≥20/40 compared to 21% of respective first eyes at baseline. Compared to non-sequentially treated fellow eyes, sequentially treated fellow eyes had higher baseline VA (p<0.001).

**Table 1:** Baseline visual acuity and visual acuity outcomes of first and fellow eyes in sequential treatment fellow eye involvement, and non-sequentially treated fellow eyes.

| Baseline characteristics<br>n = number of eyes | Sequential treatment fellow eye involvement |  |  |  | Non-sequential treatment fellow eye involvement,<br>n = 413 |
| --- | --- | --- | --- | --- | --- |
|  | First eyes<br>n = 1180 | p (paired)<br>first/fellow eyes | Fellow eyes<br>n = 1180 | p / adjusted p<br>sequential/non-sequential<br>fellow eyes |  |
| Mean VA (letters) ±SD | 54±16 | p<0.001 | 63±13 | p<0.001 | 52±16 |
| % of eyes with VA ≥ 20/40 | 21.5 | p<0.001 | 42.1 | p<0.001 | 22.3 |
| % of eyes with VA ≤20/200 | 18.1 | p<0.001 | 6.2 | p<0.001 | 23.2 |
| One-year outcomes<br>n = number of eyes | First eyes<br>n = 1094 | p (paired)<br>first/fellow eyes | Fellow eyes<br>n =961 | p / adjusted p<br>sequential/non-sequential<br>fellow eyes | Non-sequential treatment fellow eye involvement,<br>n = 321 |
| Mean VA change (letters)±SD | 5.2±15 | p<0.001 | 2.4±12 | p<0.001 | 4.1±15 |
| % of eyes with VA ≥ 20/40 | 37.9 | p<0.001 | 50.5 | p<0.001 | 34.9 |
| % of eyes with VA ≤20/200 | 17.0 | p<0.001 | 8.3 | p<0.001 | 18.7 |
| % of eyes with VA gain ≥5 letters | 52.6 | p<0.001 | 42.6 | p=0.938 | 43.0 |
| % of eyes with change in VA <15 letters | 69.4 | p<0.001 | 81.4 | p=0.006 | 74.8 |
| Mean injection number±SD | 7.7±2.1 | p=0.503 | 7.7±1.9 | p=0.646 | 7.8±2.1 |
| Two-year outcomes<br>n = number of eyes | First eyes<br>n = 1005 | p (paired)<br>first/fellow eyes | Second eyes<br>n = 781 | p / adjusted p<br>sequential/non-sequential<br>fellow eyes | Non-sequential treatment fellow eye involvement,<br>n = 255 |
| Mean VA change (letters)±SD | 5.6±15 | p<0.001 | 0.37± 14 | p<0.001 | 3.5±18 |
| % of eyes with VA≥20/40 | 38.9 | p<0.001 | 46.1 | p=0.001 | 33.7 |
| % of eyes with VA≤20/200 | 19.8 | p<0.001 | 10.4 | p=0.008 | 17.3 |
| % of eyes with VA gain ≥5 letters | 51.0 | p<0.001 | 39.2 | p=0.094 | 45.5 |
| % of eyes with change in VA <15 letters | 69.4 | p<0.001 | 81.4 | p=0.005 | 74.8 |
| Mean injection number±SD | 13±4.2 | p=0.921 | 13±3.9 | p=0.943 | 13±4 |

### Visual Acuity Outcomes

At one year, mean gain in VA was 5.2±15 letters for first eyes, 2.5±12 letters for sequentially treated fellow eyes, and 4.1±15 letters for non-sequentially treated fellow eyes (Table 1). At two years, mean gain in VA was 5.6±15 letters for first eyes and 0.65±14 letters for fellow eyes in sequentially treated patients, and 3.6±18 letters for non-sequentially treated fellow eyes. Fellow eyes showed a significantly lower gain in VA than first eyes and non-sequentially treated fellow eyes at one and two years (p <0.001). However, percentage of eyes with good vision (VA≥70 letters/>20/40) at presentation was 42%, double that of first or non-sequential fellow eyes (p<0.001) and stayed at 46% at two years, significantly higher than both other groups (p≤0.001). VA and change in VA over time is shown in Figure 2. Percentages of eyes gaining vision (change in VA ≥5 letters), stable vision (change in VA <15 letters), good vision (VA≥70 letters/>20/40), and poor vision (VA ≤35 letters/≤20/200) are shown in Table 1 and Figure 3.

**Figure 2:**
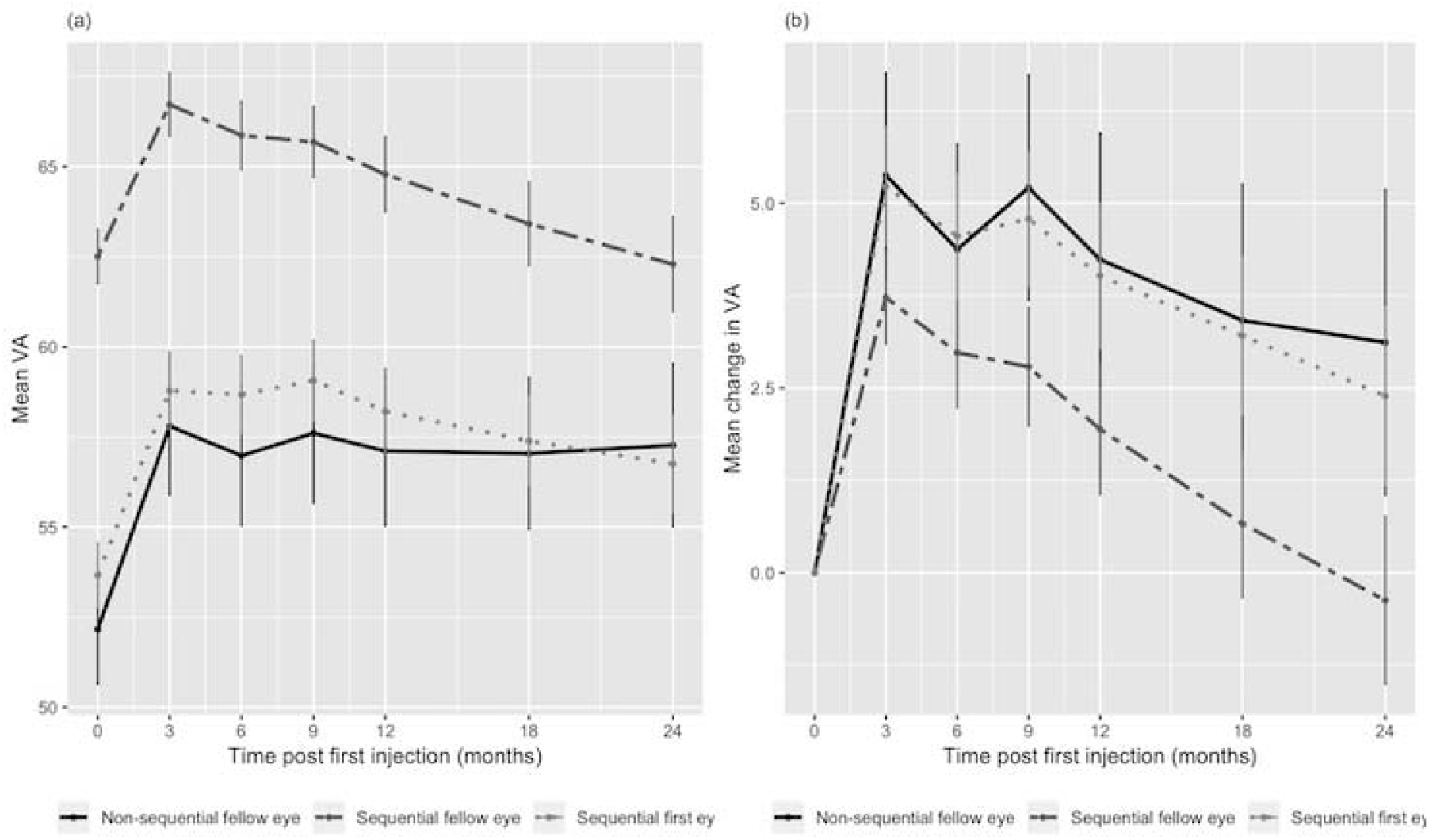
Mean visual acuity from baseline (a) and change in visual acuity (b) over time and 95% confidence interval stratified by the different groups: first and second eyes in sequentially treated fellow eye involvement and delayed-presentation fellow eyes. VA - visual acuity

**Figure 3.**
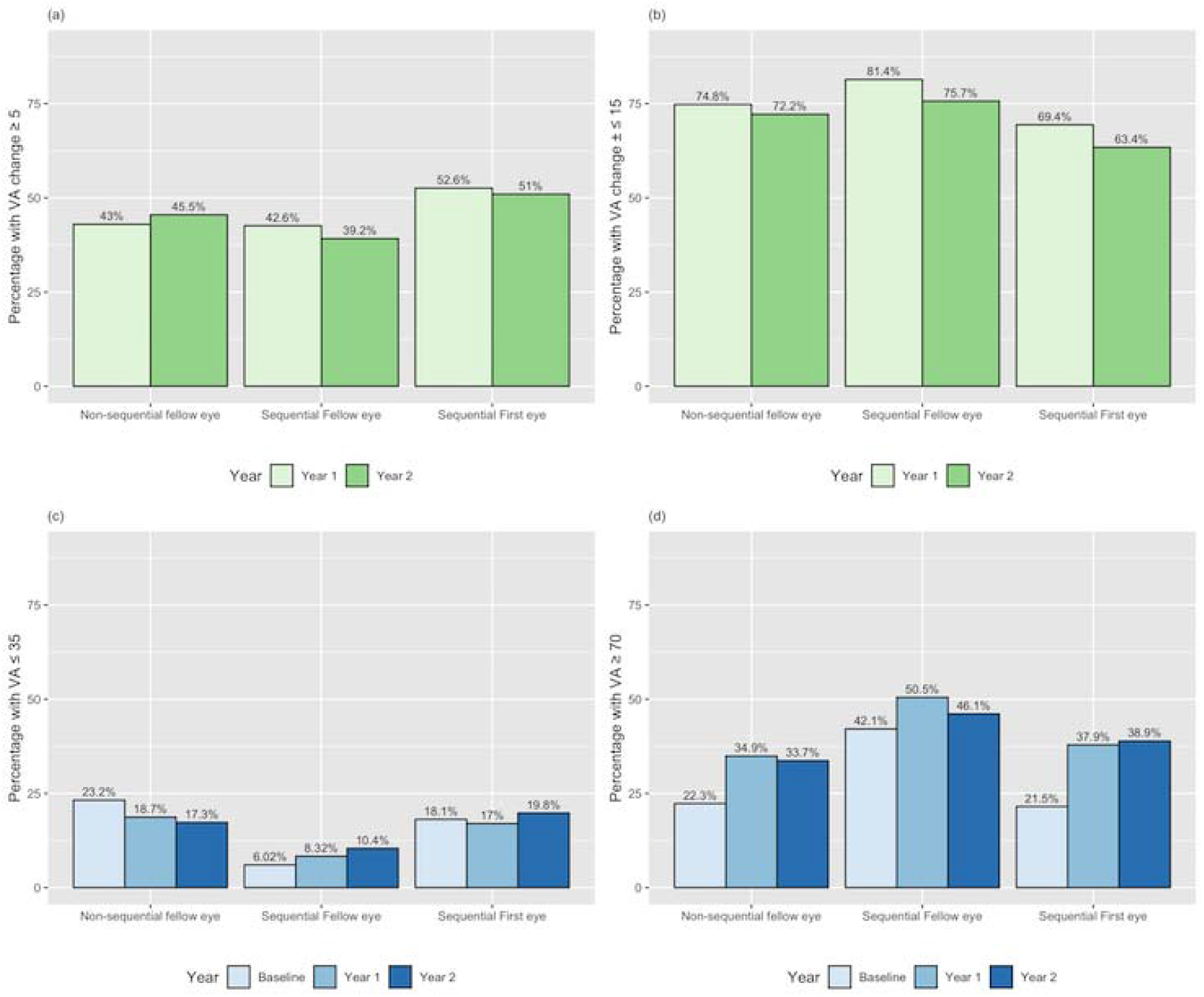
a: Percentages of eyes with gain in VA (≥ 5 letters) at one and two years for fellow eyes in sequential/non-sequential treatment, and first eyes in sequential treatment. b: Percentages of eyes with stable vision (change in VA ≤±14 letters) for fellow eyes in sequential/non-sequential treatment, and first eyes in sequential treatment. c: Percentages of eyes with poor vision (VA≤ 35 letters or 20/200) for fellow eyes in sequential/non-sequential treatment, and first eyes in sequential treatment. d: Percentages of eyes with good vision (VA≥ 70 letters or 20/40) for fellow eyes in sequential/non-sequential treatment, and first eyes in sequential treatment. VA - visual acuity

### Time to Involvement of Second Eye

Median time interval between involvement of first and fellow eye in sequential involvement was 71 weeks (interquartile range: 27-147 weeks). Chance of involvement of fellow eye involvement for eyes starting treatment in one eye 21% (486 eyes) at one year and 32% (742 eyes) at two years, and it was dependent on age at presentation of the first eye: At two years, the risk of fellow eye involvement was 20% for patients younger than 60 years and 40% for patients in their eighties. Survival analysis of fellow-eye involvement is shown in Figure 4.

**Figure 4.**
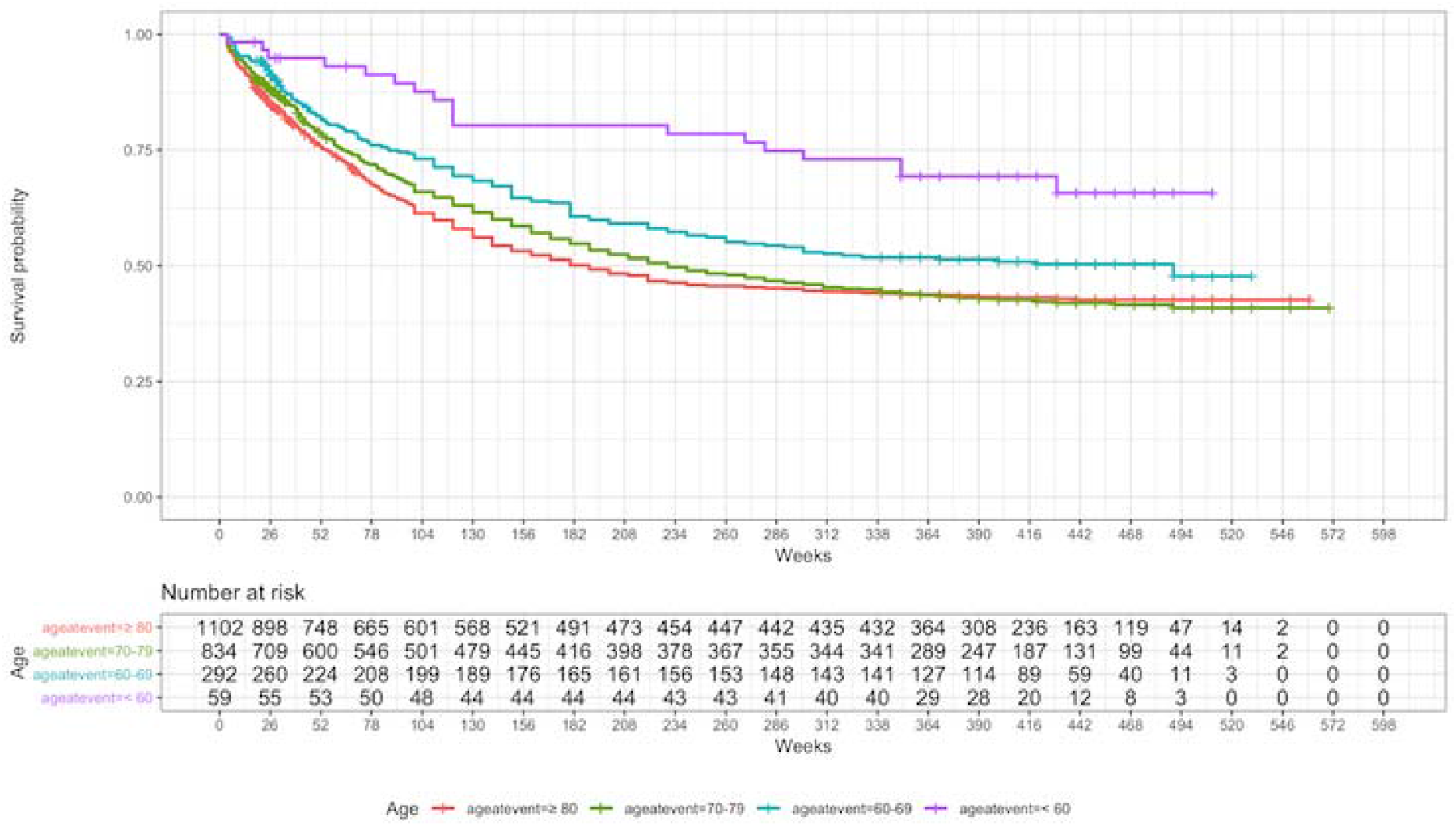
Survival probability for fellow eye involvement over time (weeks).

### Injection Frequency

Mean number of injections was 8 in all groups at one year and 13 at two years with no significant differences between the groups (Table 1).

## DISCUSSION

Our study shows that fellow eye involvement of nAMD affects 20 to 40% over a two-year period, depending on age at presentation of the first eye, and that there is a significant difference in both baseline VA and VA outcomes depending on characteristics of the first eye.

Fellow eye involvement in nAMD is very common, reaching 20-40% depending on age at presentation after two years in our cohort. This rate falls within the range reported in the Comparison of AMD Treatments Trials (20.6% at two years) and other studies.[12,13,23–25] With demographic ageing, sight loss and blindness is predicted to increase by 2.4% from 2013 until 2050, reaching approximately 4 million people in the UK and the share of AMD is estimated to rise from 23% to 30%, representing 1.23 million people.[8] AMD is thus a major and growing contributor to healthcare burden. Considering the annual societal cost per bilaterally treated AMD patient estimated at 5300€ in 2005, the frequency of bilateral involvement makes this patient cohort an important target for vision loss prevention and healthcare cost reduction.[7]

In sequential treatment, fellow eyes have a higher baseline VA and maintain good vision over two years of treatment despite the absence of an initial gain in VA comparable to first eyes. This ceiling effect has been well-described and implies a rationale for earliest possible detection and treatment of neovascular changes in AMD.[13,16] Example A (Figure 1) reflects this fellow eye advantage, in which neovascular AMD in the fellow eye was detected pre-symptomatically and treatment was started immediately. Patients might profit from the routinely performed bilateral OCT imaging at every visit, be more vigilant of VA changes in their fellow eye while undergoing treatment for the first eye, and profit from the already in-place pathway to access treatment for the fellow eye quickly. This effect has led to a discussion about strategies for early detection of nAMD and optimal interval of monitoring of AMD patients.[12,13,26] Specifically analysis of imaging biomarkers, possibly aided by artificial intelligence, might prove to be key in risk stratification of fellow eye involvement.[24,27,28]

Our study demonstrates that non-sequentially treated fellow eyes do not share the typical fellow eye characteristics. They start treatment with a relatively low baseline VA and their gain in VA is higher, very similar to first eyes of sequentially treated patients. Explanations for this could be that patients with non-treatable advanced neovascular disease in the first eye are not regularly monitored or that there is systematic delay in access to treatment. This is supported by the existing lack of awareness of AMD and evidence of substantial delay from symptoms to treatment in the UK AMD care pathways.[29] Interestingly, in this cohort of patients, vision loss secondary to macular scarring in the first eye does not appear to result in increased vigilance that could lead to early detection of fellow eye involvement. One might argue that scarring in the first eye implies more aggressive disease causing worse VA at presentation of fellow eyes, but the similar VA gain over time to first eyes in our cohort does not support this theory. To our knowledge, the findings on non-sequential fellow eye involvement have not been reported before and highlight the arguably most vulnerable cohort of patients in which vision loss in their fellow and better or functionally only seeing eye will lead to significant visual impairment and socioeconomic burden.[7,8]

The limitations of this study lie within its retrospective nature based on EMR and the LTFU. However, comparisons of baseline characteristics based on the complete cohorts are unaffected by LTFU and it has been shown that changes in VA are comparable in cohorts of different follow-up periods.[16] Strengths of this study include the large sample size for fellow eye cohorts and the the quality of data coming from one single center and a curated database with additional substantial manual cleaning. Additionally and maybe most importantly, we encourage an open science approach to replicate our results with freely available depersonalized raw data and code.

In conclusion, this study highlights the superior visual outcomes of fellow eyes compared to first eyes in the common scenario of sequential fellow eye involvement in nAMD as well as the inferior outcomes of fellow eyes in case of untreated late stage neovascular disease in the first eye. Future research should account for those idiosyncratic subgroups of fellow eyes undergoing treatment for nAMD, as these could prove to represent the span of quality of care in AMD treatment.

## Supporting information

Supplementary sFigure1

## Abbreviations

VEGF: - vascular endothelial growth factor
AMD: - age related macular degeneration
nAMD: - neovascular age related macular degeneration
VA: - visual acuity
UK: - United Kingdom
EMR: - electronic medical record
ETDRS: - Early Treatment Diabetic Retinopathy Study
ICHOM: - International Consortium for Health Outcomes Measurement
RCTs: - randomized controlled trials

