## Supplementary sFigure1 for "The Moorfields AMD Database Report 2 - Fellow Eye Involvement with Neovascular Age-related Macular Degeneration"

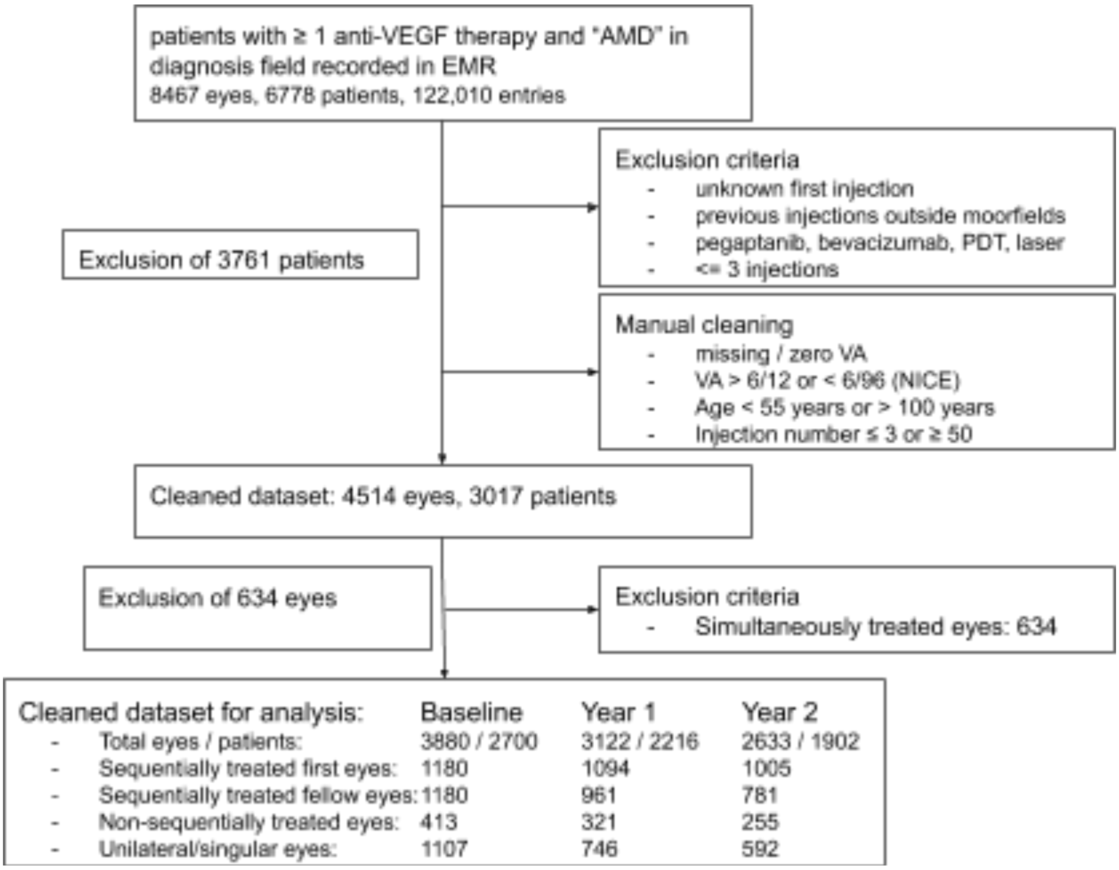


**Supplementary sFigure 1:** Consolidated Standards of Reporting Trials-style diagram showing patients / eyes included in the analysis

VEGF- vascular endothelial growth factor, AMD - age related macular degeneration, PDT - photodynamic therapy, VA - visual acuity, NICE national institute for health and care excellence
